# Kynurenine and NAD+ Pathways are Associated with Macrophage Content and Polarization in Carotid Plaques

**DOI:** 10.1101/2025.04.14.648852

**Authors:** Charlotte J. Teunis, Lobke F. Zijlstra, Johannes H.M. Levels, Kasper T. Vinten, Maria M. Trętowicz, Barend Mol, Judith C. Sluimer, Michal Mokry, Andrew J. Murphy, Jeffrey Kroon, Erik Stroes, Dominique P.V. de Kleijn, Riekelt H. Houtkooper, Annette E. Neele, Nordin M.J. Hanssen

**Affiliations:** Amsterdam UMC, location University of Amsterdam, Department of Vascular Medicine, Meibergdreef 9, Amsterdam, the Netherlands; Amsterdam UMC, location University of Amsterdam, Department of Medical Biochemistry, Meibergdreef 9, Amsterdam, the Netherlands; Amsterdam Cardiovascular Sciences, Atherosclerosis & Ischemic Syndromes, Amsterdam, the Netherlands; Amsterdam Institute for Immunology and Infectious disease, Amsterdam, the Netherlands; Amsterdam UMC location University of Amsterdam, Department of Experimental Vascular Medicine, Meibergdreef 9, Amsterdam, Netherlands; Laboratory Genetic Metabolic Diseases, Amsterdam UMC, University of Amsterdam, the Netherlands; Amsterdam Gastroenterology, Endocrinology, and Metabolism Institute, Amsterdam, the Netherlands; Department of Vascular Surgery, University Medical Center Utrecht, Utrecht University, the Netherlands; Department of Pathology, Cardiovascular Research Institute Maastricht (CARIM), Maastricht University Medical Center (MUMC), Maastricht, the Netherlands; Department of Medicine 2 (Nephrology, Clinical Immunology, Rheumatology, Hypertension), RWTH Aachen University, Medical Faculty, Aachen, Germany; BHF Centre for Cardiovascular Sciences, University of Edinburgh, Edinburgh, UK; Laboratory of Experimental Cardiology University Medical Center Utrecht, Utrecht University, the Netherlands; Central Diagnostic Laboratory, University Medical Center Utrecht, Utrecht University, the Netherlands; Haematopoiesis and Leukocyte Biology, Baker IDI Heart and Diabetes Institute, Melbourne, 3004, Australia; Department of Immunology, Monash University, Melbourne, 3004, Australia; Laboratory of Angiogenesis and Vascular Metabolism, VIB-KU Leuven Center for Cancer Biology, VIB, Belgium; Laboratory of Angiogenesis and Vascular Metabolism, Department of Oncology, KU Leuven and Leuven Cancer Institute (LKI), Belgium; Diabeter Centrum Amsterdam, Amsterdam, the Netherlands; Emma Center for Personalized Medicine, Amsterdam UMC, Amsterdam, the Netherlands

**Author notes:** Corresponding author:* Nordin M.J. Hanssen, *Email:.

## Abstract

**Background and Aims:** Metabolism dictates macrophage function and plays a central role in atherosclerotic plaque progression. The kynurenine pathway, which metabolizes the majority of the essential amino acid tryptophan, plays a pivotal role in regulating immune responses and supporting NAD+ synthesis, essential for cellular energy metabolism. Higher circulating kynurenine levels are associated with cardiovascular disease, yet their role in atherosclerotic plaques is unclear. This study aims to investigate the underlying mechanisms driving increased kynurenine concentrations in plaques and to determine whether kynurenine serves as a mere biomarker of low-grade inflammation or reflects specific macrophage-driven metabolic alterations that could position it as a potential therapeutic target.

**Methods:** We used histological and transcriptomic data from two biobanks: the Athero Express Biobank (AE; n=91) and Maastricht human plaque study (MaasHPS, n= 26). Macrophages were identified through CD68 staining in AE, and M1/M2-like macrophage subtypes were distinguished by iNOS/CD68 and arginase/CD68 expression in MAASHPS. Primary human monocyte-derived cultured macrophages were polarized into M1- and M2-like phenotypes for using IFN-γ and IL-4, respectively. Tryptophan, kynurenine and/or NAD+ concentrations in plaques were quantified usingliquid chromatography and metabolomics analyses.

**Results:** Kynurenine concentrations were significantly higher in plaques with greater macrophage density (p = 0.023). Transcriptomic analysis in AE revealed upregulation of *IDO2, AFMID*, and *KYNU* in plaques with increased macrophage infiltration (p < 0.05), but not *IDO1* (p = 0.16). In the MAASHPS biobank, higher *IDO1, KYNU*, and *KMO* expression correlated negatively with M2 marker positive macrophages (p < 0.001), while *HAAO* correlated positively (p < 0.01). In vitro, M1-like macrophages showed increased *IDO1* and reduced *QPRT* expression compared to M2-like macrophages. We found that this disruption in kynurenine pathway gene expression led to decreased NAD+ concentrations in M1-like macrophages compared to M2-like macrophages in vitro.

**Conclusion:** Higher kynurenine levels in atherosclerotic plaques are increased by the increased presence of M1 macrophages, likely driven by both an increased IDO1 activity and reduced *QPRT* gene expression. This leads to decreased concentrations of NAD+, potentially determining the phenotype of the macrophages. Future studies should address whether modulation of the kynurenine pathway restores NAD+ metabolism and leads to a decrease in inflammation and an increased stable plaque phenotype.

## Introduction

Macrophages are central to the inflammatory and immune processes that drive atherosclerosis (1). Within the atherosclerotic plaque, macrophages represent a heterogeneous population that dynamically adapts to their environment, influenced by oxidized lipids, inflammatory triggers, and other local factors. Consequently, dysregulated macrophage activation, is a key orchestrator of atherosclerosis progression (2, 3). Inflammatory “M1” macrophages, are associated with plaque progression and instability, whereas anti-inflammatory “M2” macrophages are linked to plaque stability and resolution. Together they represent two extreme ends of the macrophage activation spectrum (4, 5). Targeting the metabolic pathways that underpin macrophage activation and function could provide novel therapeutic approaches for reducing inflammation and improving plaque stability. Kynurenine metabolites, formed from the essential amino acid tryptophan (Trp) (6), have been implicated in the pathogenesis of cardiovascular disease (CVD), with higher circulating kynurenine levels repeatedly associated with increased cardiovascular risk (7-9). Despite these associations, it remains unclear whether kynurenine metabolites function as biomarkers, active mediators of disease, or both. (10-12). Elucidating their role in CVD could unveil therapeutic opportunities targeting this pathway, because inflammation as the hallmark of atherosclerosis in turn is a key driver of kynurenine pathway activation (13, 14). Indeed, inflammatory cytokines such as interleukin-1 (IL-1), interleukin-6 (IL-6), and interferon-gamma (IFN-γ) potently induce indoleamine 2,3-dioxygenase 1 (*IDO1*), which catalyzes the rate-limiting conversion of Trp to kynurenine (15, 16). In atherosclerotic models, *IDO1* expression is predominantly observed in macrophages, and IDO1 activity correlates with kynurenine accumulation (17). However, the functional significance of this pathway in atherosclerosis is debated. While some rodent studies have linked *IDO1* activation to atherogenesis, others suggest a protective role (18-20). Beyond IDO1, downstream metabolites such as kynurenine and nicotinamide adenine dinucleotide (NAD+) are known to influence cellular metabolism. Yet, their specific contributions to inflammation, immune cell function, and plaque progression remain poorly understood. Whether higher kynurenine levels in plaques reflect macrophage abundance, macrophage phenotype, or broader metabolic changes remains to be determined.

NAD+, a central cofactor in cellular energy metabolism, has emerged as a potential link between the kynurenine pathway and macrophage function (21, 22). Differentiated macrophage subsets exhibit distinct metabolic profiles, with M1-like macrophages predominantly relying on glycolysis, while M2-like macrophages favor oxidative phosphorylation (23, 24). This metabolic divergence extends to NAD+ biosynthesis, with kynurenine pathway activation and de novo NAD+ synthesis being elevated in M1-like macrophages (21). The enzyme quinolinic acid phosphoribosyltransferase (QPRT) catalyzes the conversion of quinolinic acid, the kynurenine pathway’s terminal metabolite, into nicotinic acid mononucleotide, a precursor of NAD+ (25). Disruptions in this pathway could therefore have profound effects on macrophage function and plaque biology.

In light of these considerations, the objective of this study was to examine how alterations in the kynurenine pathway, driven by IDO1 activity, affect macrophage function within atherosclerotic plaques. By integrating metabolomic and transcriptomic data, we sought to unravel the link between kynurenine accumulation, disruptions in NAD+ synthesis, and macrophage-driven inflammation.

## Methods

### Study Design and Biobanks

This study utilized data from two independent biobanks: the Athero Express Biobank (AE), an ongoing prospective biobank (26) and the Maastricht human plaque study (MaasHPS) biobank. Both biobanks consisted of atherosclerotic plaques obtained from patients undergoing carotid endarterectomy. The AE biobank included 117 samples and in the MAASHPS biobank plaque samples were obtained from 23 patients. For each patient, paired plaque segments were collected, resulting in a total of 46 segments.

The study received ethical approval from the respective institutional review boards of each biobank in accordance with the Declaration of Helsinki.

### Histological staining

In the AE biobank, macrophage content in atherosclerotic plaques was assessed through immunohistochemical staining for CD68. Macrophages were classified into four groups based on staining intensity: no staining, minor staining, moderate staining, and heavy staining. These classifications were used to stratify plaques for subsequent analyses. Within the MAASHPS biobank, macrophage phenotypes within the plaques were also characterized using immunohistochemistry, where iNOS was used as a marker for pro-inflammatory M1 macrophages, while ARG1 was used for anti-inflammatory M2 macrophages. The relative area of M1 and M2 macrophages was quantified as a percentage of total CD68+ macrophage areas in the atherosclerotic lesion (27).

### HPLC Metabolomics Analysis

Targeted metabolomic analysis was performed to quantify the concentrations of tryptophan and kynurenine in plaque lysate from the Athero Express biobank using an in-house HPLC method. A 100 µl sample of atheroma lysate was spiked with Nitro-Tyrosine (Sigma-Aldrich) as an internal standard and subsequently precipitated by the addition of 55.6 µl with perchloric Acid (10%) (Sigma-Aldrich). All samples were vortexed for 10 sec, incubated on ice for 15 min and finally spun for 10 min in an Eppendorf centrifuge at maximum speed. A 50 µl aliquot of the supernatant was transferred into an HPLC vial for further analysis. All samples and standards were stored in a Jasco autosampler (AS4285, Jasco Benelux) at 10 °C. A 10 μl aliquot was injected into the HPLC system for all standards (tryptophan and Kynurenine, Sigma Aldrich) or samples. Metabolite separation was carried out on a Microsphere C18 column (100 mm × 4.6 mm, dp: 3 μm, Agilent Technologies). The column temperature was maintained at 35 °C and variable flow conditions (0.8-1.5 mLl/min) were applied for the mobile phase (NaAc,15 mM, pH 5.5)/acetonitrile mix using a Jasco quaternary pump (PU4285, Jasco Benelux) during each run. The applied acetonitrile gradient was varied from 1% −20% and detection was done at 270 nm with a Jasco UV detector at 30 °C (UV4075, Jasco Benelux). Final data interpretation and quantification were performed using Chrom-Nav (v 2.0). Samples that yielded no results for either metabolite were excluded from the analysis. Finally, the metabolite concentrations were normalized to total protein concentration.

### LC-MS Metabolomics Analysis

For metabolomics analysis of the in vitro cultured macrophages, cells were detached with TrypleE (Gibco) washed with 0,9% NaCl and metabolically quenched with 500 μl ice-cold methanol (LiChrosolv®). Metabolomics was performed as described previously, with minor adjustments (28, 29). The following amounts of internal standard dissolved in Milli-Q water were added to each sample: adenosine-^15^N_5_-monophosphate (5 nmol), adenosine-^15^N_5_-triphosphate (5 nmol), D_4_-alanine (0.5 nmol), D_7_-arginine (0.5 nmol), D_3_-aspartic acid (0.5 nmol), D_3_-carnitine (0.5 nmol), D_4_-citric acid (0.5 nmol), ^13^C_1_-citrulline (0.5 nmol), ^13^C_6_-fructose-1,6-diphosphate (1 nmol), ^13^C_2_-glycine (5 nmol), guanosine-^15^N_5_-monophosphate (5 nmol), guanosine-^15^N_5_-triphosphate (5 nmol), ^13^C_6_-glucose (10 nmol), ^13^C_6_-glucose-6-phosphate (1 nmol), D_3_-glutamic acid (0.5 nmol), D_5_-glutamine (0.5 nmol), D_5_-glutathione (1 nmol), ^13^C_6_-isoleucine (0.5 nmol), D_3_-lactic acid (1 nmol), D_3_-leucine (0.5 nmol), D_4_-lysine (0.5 nmol), D_3_-methionine (0.5 nmol), D_6_-ornithine (0.5 nmol), D_5_-phenylalanine (0.5 nmol), D_7_-proline (0.5 nmol), ^13^C_3_-pyruvate (0.5 nmol), D_3_-serine (0.5 nmol), D_6_-succinic acid (0.5 nmol), D_4_-thymine (1 nmol), D_5_-tryptophan (0.5 nmol), D_4_-tyrosine (0.5 nmol), D_8_-valine (0.5 nmol), ^13^C_5_-NAD+ (1.25nmol). The samples were adjusted to a total volume of 1 mL with Milli-Q water. 1 mL of chloroform was added to each tube and samples were thoroughly mixed, before centrifugation for 10 min at 14.000 rpm to facilitate layer separation. The top layer, containing the polar phase, was transferred to a clean 1.5 mL tube and dried using a vacuum concentrator at 90°C. The remaining portion was dried at room temperature, and protein content was determined using the Thermo Scientific™ Pierce™ BCA Protein Assay kit (#23225) following the manufacturer’s instructions. Dried samples were reconstituted in 100 μL 6:4 (v/v) methanol:water. Metabolites were analyzed using a Waters Acquity ultra-high performance liquid chromatography system coupled to a Bruker Impact II™ Ultra-High Resolution Qq-Time-Of-Flight mass spectrometer. Samples were kept at 15°C during analysis and 5 μL of each sample was injected. Chromatographic separation was achieved using a Waters Atlantis Premier BEH Z-HILIC column (PEEK 100 x 2.1 mm, 1.7 μm particle size). Column temperature was held at 30°C. Mobile phase consisted of (A) 1:9 (v/v) acetonitrile:ammonium bicarbonate (15mM in water) and (B) 9:1 (v/v) acetonitrile:ammonium bicarbonate (15mM in water). Using a flow rate of 0.400 mL/min, the LC gradient consisted of: Dwell at 95% Solvent B, 0-2 min; Ramp to 60% Solvent B at 2-11 min; Dwell at 60% Solvent B, 11-12 min; Ramp to 95% B at 12-13 min; Dwell at 95% Solvent B, 13-15 min. MS data was acquired using negative and positive ionization in full scan mode over the range of m/z 50-1200. Data was analyzed using Bruker TASQ software version 2.1.22.3. All reported metabolite intensities were normalized to protein content, as well as to internal standards with comparable retention times and response in the MS. Metabolite identification has been based on a combination of accurate mass, (relative) retention times, ion mobility data and fragmentation spectra, compared to the analysis of a library of standards.

### Transcriptomic Analysis

Detailed descriptions of the transcriptomic methods used in both the AE biobank (30) and MaasHPS biobank (27) are available in previous publications.

Briefly, in the AE biobank, atherosclerotic plaques were ground in liquid nitrogen, homogenized with Tripure, and RNA was isolated via chloroform extraction and isopropanol precipitation. RNA quality was assessed, and the CEL-seq2 method was chosen for library preparation due to its superior read mappability. This method employed unique molecular identifiers (UMIs) for precise RNA molecule quantification. Libraries were cleaned using AMPure XP beads and sequenced on the Illumina NextSeq500 platform (paired-end, 2 × 75 bp).

In the MAASHPS biobank, RNA was extracted from snap-frozen plaque segments using guanidinium thiocyanate and further purified with the Nucleospin RNA II kit (Macherey-Nagel). RNA integrity was verified using an Agilent 2100 Bioanalyzer, and samples with a RIN >6.0 were included. Biotinylated cRNA was synthesized and hybridized to Illumina HumanRef-8 v2.0 expression BeadChips for transcriptomic profiling. Preprocessing and normalization were performed using variance stabilizing transformation and robust spline normalization.

### Cell culture

Peripheral blood mononuclear cells (PBMCs) were isolated from healthy controls (50% male/ 50% female) from buffy coats (Sanquin blood supply, Amsterdam, the Netherlands) through density centrifugation using Lymphoprep™ (Axis-Shield). Written informed consent was obtained from all donors in accordance with the principles of the Declaration of Helsinki (2013). CD14^+^ monocytes were isolated using CD14 magnetic beads and MACS® cell separation columns (Miltenyi). Monocytes were plated in 24-well tissue culture plates at a density of 500.000 cells per well and differentiated into macrophages for 7 days using 50 ng/mL of human M-CSF (Miltenyi) and cultured in glucose free Roswell Park Memorial Institute (RPMI) 1640 Medium supplemented with 10% heat-inactivated Fetal Calf Serum (FCS), 100 U/mL Penicillin, 100ug/mL Streptomycin, 5 mM Glucose Solution (Gibco™), and 25 mM HEPES (Gibco™). On day 3 and day 6, all medium was removed and substituted by fresh RPMI with 10% FCS and 50ng/mL M-CSF. On day 7, macrophages were stimulated for 24h with either 50 ng/mL IFN-γ (R&D systems), 50 ng/mL IL-4 (PeproTech) or left unstimulated as control to induce an M1-like or a M2-like phenotype.

### RNA isolation and qPCR of in-vitro macrophages

Total RNA was isolated from cells using the Thermo Scientific GeneJET RNA Purification Kit (Thermo Fisher) following manufacturers protocol. Obtained RNA was reverse transcribed into cDNA with use of the High-Capacity cDNA Reverse Transcription Kit (Applied Biosystems™). Fast SYBR™ (Applied Biosystems™) was used for real-time quantitative qPCR (qPCR), performed on a QuantStudio 5 real time PCR (Applied Biosystems™). mRNA expression levels were normalized tot the average of the two housekeeping genes *HRPT* and *GNB21L*. Primer sequences are listed in Supplementary Table 1

#### Statistical Analysis

The baseline characteristics were presented as mean ± standard deviation (SD) for parametric variables, median [interquartile range (IQR)] for non-parametric variables, and counts (percentages) for categorical variables. Missing data in categorical variables were included in percentage calculations but excluded from statistical comparisons. The differences between groups were assessed using ANOVA for parametric variables, the Kruskal-Wallis test for non-parametric variables, and chi-square tests for categorical variables.

To examine any trends across the macrophage staining groups, the Jonckheere-Terpstra test was employed, with predefined hypotheses tested for each metabolite and inflammatory marker.

All statistical analyses were performed using R (version 4.2.1) with a p-value <0.05 considered statistically significant.

### Results

In the AE biobank, 91 out of 117 samples yielded enough material for targeted metabolomic analysis. Baseline characteristics of this biobank are shown in Table 1. The mean age was 67.6 ± 9.1 years and did not differ significantly across macrophage staining groups (p = 0.8). The proportion of female participants varied significantly, with the lowest percentage in the heavy staining group (p = 0.03). There was a significant difference in the prevalence of type 2 diabetes mellitus among groups, being highest in the minor staining group (p = 0.001). Statin use was more prevalent in the moderate and heavy staining groups compared to the no staining group (p = 0.003). No significant differences were observed in other characteristics, i.e. BMI, eGFR, smoking status, and anti-hypertensive medication use, between the groups.

**Table 1.** Baseline characteristics of the AtheroExpress biobank.

| Variable | Macrophage Staining |  |  |  |  | P-value |
| --- | --- | --- | --- | --- | --- | --- |
|  | All<br>(n = 91) | No staining<br>(n = 21) | Minor<br>staining<br>(n = 19) | Moderate<br>staining<br>(n = 30) | Heavy staining<br>(n = 21) |  |
| Age | 67.6 ± 9.1 | 70.7 ± 6.5 | 63.6 ± 9.2 | 70.0 ± 8.8 | 69 ± 10.7 | 0.835 |
| Female | 25 (27.5%) | 9 (42.9%) | 8 (42.1%) | 6 (20%) | 2 (9.5%) | **0.033** |
| Smoker | 37 (40.7%) | 9 (42.9%) | 9 (47.4%) | 11 (36.7%) | 8 (38.1%) | 0.885 |
| DM | 13 (14.3%) | 1 (4.8%) | 8 (42.1%) | 2 (6.7%) | 2 (9.5%) | **0.004** |
| BMI | 25.9 [24.1 -<br>28.4] | 26.2 [24.1 -<br>28.4] | 27.4 [25.3<br>- 29.4] | 25.1 [24.0 - 27.9] | 25.2 [23.7 - 26.4] | 0.291 |
| eGFR | 70.7 ± 18.2 | 69.2 ± 14.6 | 76.6 ± 20.3 | 69.6 ± 18.4 | 68.2 ± 19.2 | 0.573 |
| Statins | 64 (70.3%) | 9 (42.9%) | 12 (63.2%) | 27 (90%) | 16 (76.2%) | **0.003** |
| AHD | 67 (73.6%) | 16 (76.2%) | 11 (57.9%) | 23 (76.7%) | 17 (81%) | 0.592 |

Data are presented as the mean ± SD for parametric variables and as the median [IQR] for non-parametric variables. Trends across parametric variables (age, eGFR, BMI)
were compared across macrophage staining groups using linear regression. Categorical variables are shown as counts (%) and were analyzed using the chi-square test.
Abbreviations: AHD: anti-hypertensive drugs, DM; diabetes mellitus type II, TC; total cholesterol, TG; triglycerides.

### Kynurenine Accumulates in Plaques with Higher Macrophage Content, Without a Shift in the [Kyn]/[Trp] Ratio

The distribution of kynurenine and tryptophan as well as their ratio, was investigated across the atherosclerotic plaques stratified by macrophage staining intensity (Figure 1). The concentration of kynurenine was higher with increasing macrophage staining intensity (p = 0.023, Figure 1). In contrast, tryptophan levels did not exhibit a significant trend across the macrophage staining groups (p = 0.999, figure 1). The kynurenine-to-tryptophan ratio, which is commonly used as an indirect marker of indoleamine 2,3-dioxygenase (IDO) activity, also demonstrated no significant trend over different macrophage staining groups (p = 0.989) implying that IDO activity is not merely a reflection of macrophage content in the atherosclerotic plaque.

**Figure 1.**
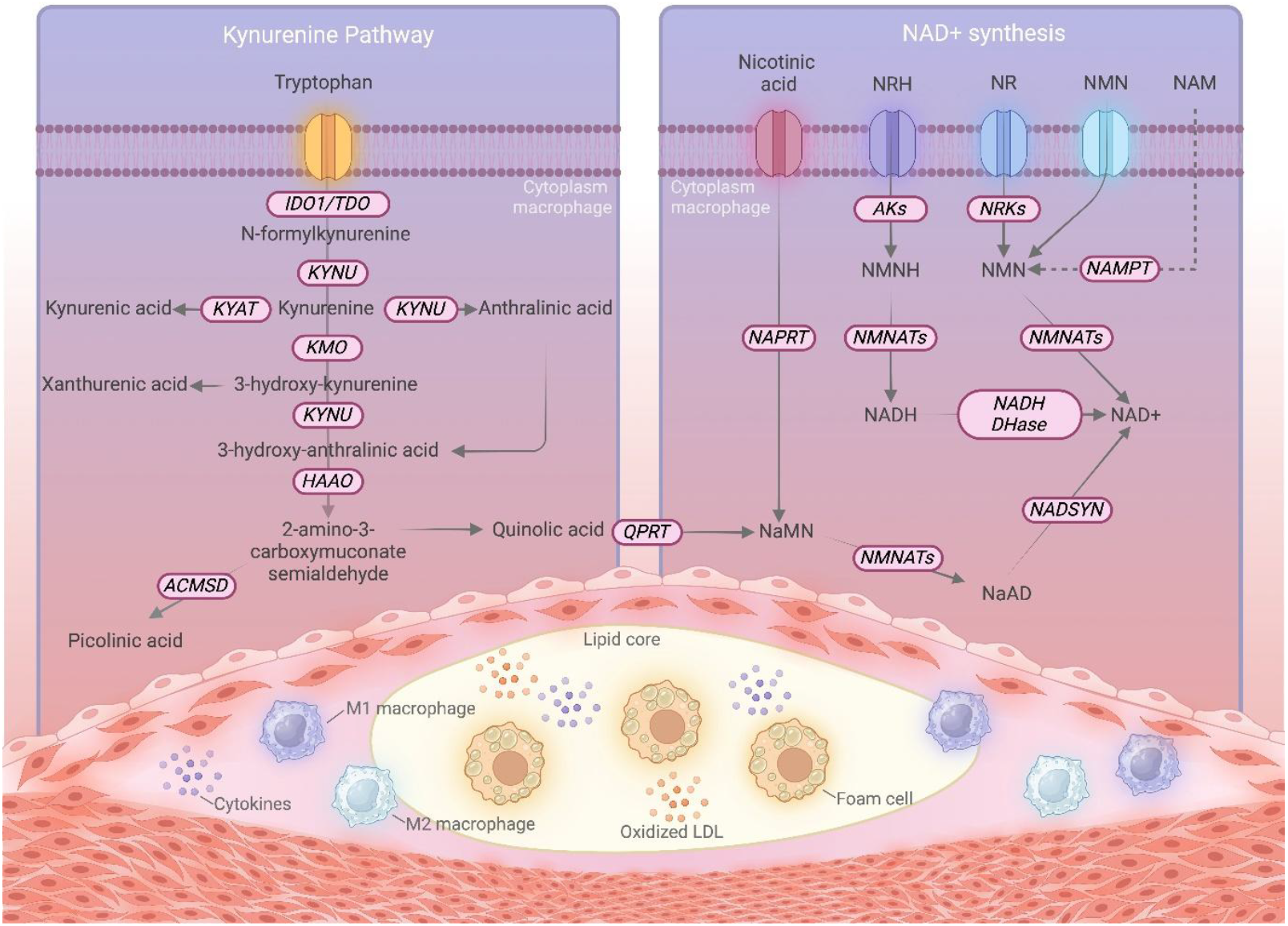
Kynurenine and NAD+ synthesis in atherosclerosis. Abbreviations: ACMSD – Aminocarboxymuconate Semialdehyde Decarboxylase, Aks – Aryl Kynurenine Synthase, HAAO – 3-Hydroxyanthranilate 3,4-Dioxygenase, IDO1 – Indoleamine 2,3-Dioxygenase 1, KMO – Kynurenine 3-Monooxygenase, KYAT – Kynurenine Aminotransferase, KYNU – Kynureninase, NADH DHase – NADH Dehydrogenase, NADSYN – NAD+ Synthetase, NAM – Nicotinamide, NAMPT – Nicotinamide Phosphoribosyltransferase, NAPRT – Nicotinic Acid Phosphoribosyltransferase, NMN – Nicotinamide Mononucleotide, NMNATs – Nicotinamide Mononucleotide Adenylyltransferases, NR – Nicotinamide Riboside, NRH – Reduced Nicotinamide Riboside, NRKs – Nicotinamide Riboside Kinases, QPRT – Quinolinate Phosphoribosyltransferase, TDO – Tryptophan 2,3-Dioxygenase.

**Figure 2.**
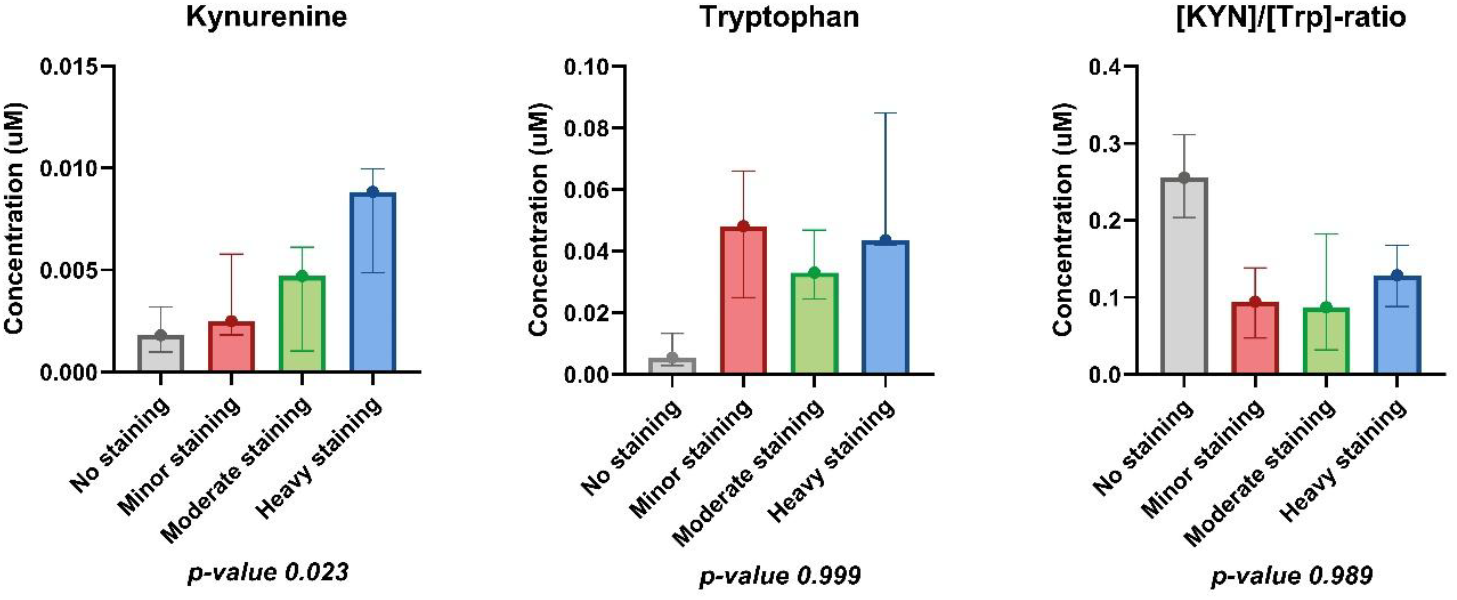
Box plots showing the distribution of log-transformed tryptophan (μM), kynurenine (μM), and kynurenine-to-tryptophan ratio in atherosclerotic plaques with increasing macrophage staining intensity. Boxes represent the median and interquartile range (IQR), whiskers extend to 1.5× IQR.. Trends across groups were assessed using the Jonckheere-Terpstra test.

### Kynurenine Pathway Gene Expression Varies with Macrophage Burden in Atherosclerotic Plaques

The subsequent objective was to utilize transcriptomics analysis to ascertain the potential pathways associated with the formation and metabolism of kynurenine and the macrophage burden of the plaque. Transcriptomic analysis of genes in the AE biobank plaques revealed significant differences in the expression of *IDO2* (p < 0.05), *AFMID* (p < 0.05), and *KYNU* (p < 0.01) between plaques with increasing macrophage infiltration (Figure 3). No statistically significant differences were observed in *IDO1* gene expression between these groups (p = 0.26). No significant differences were observed in the gene expression of kynurenine pathway enzymes between male and female or across other plaque characteristics, including smooth muscle cell content, intraplaque hemorrhage, calcification, collagen content, or plaque phenotype. Among the genes involved in other parts of the NAD pathway than the de novo NAD+ synthesis, only *NADSYN1* and *NAMPT* genes expression from the salvage pathway showed a statistically significant difference across macrophage groups (p < 0.05 and p < 0.001; Figure 3A).

**Figure 3.**
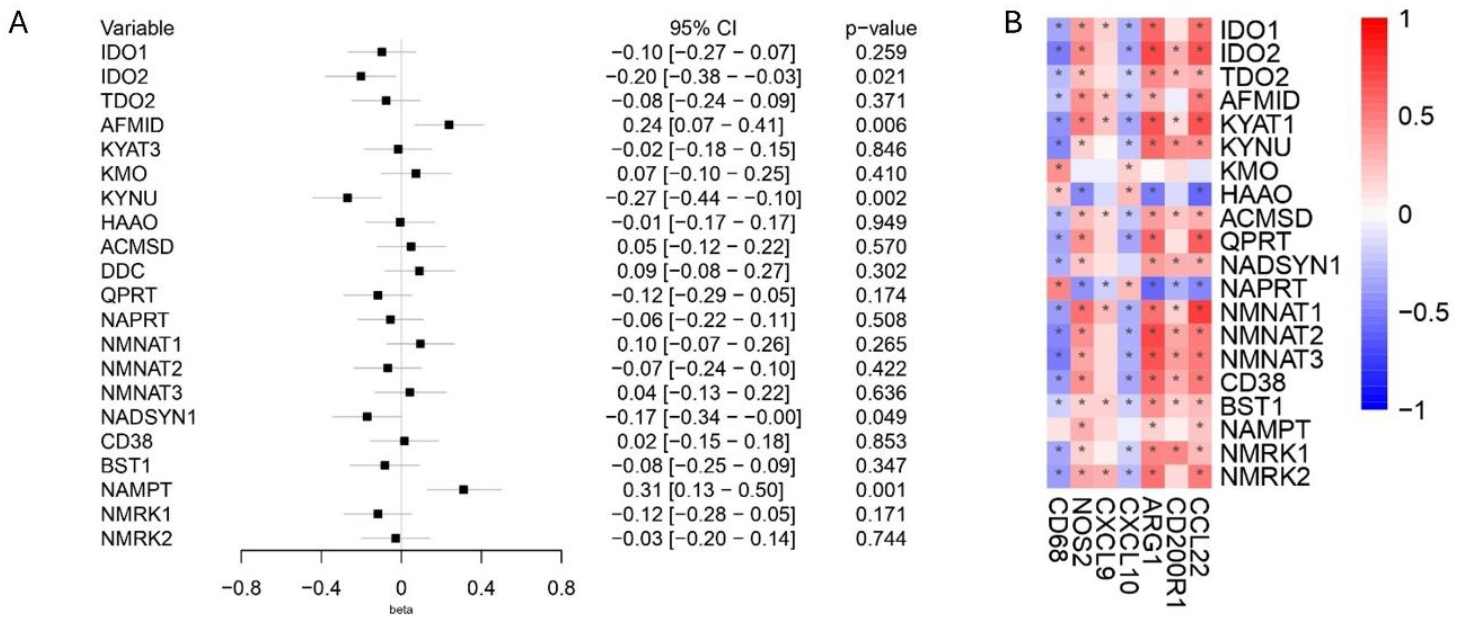
A. Forest plot showing the β-coefficient for the association between the presence of histologically scored macrophages in advanced carotid lesions and the expression levels of various genes. Error bars represent 95% confidence intervals. B. Heatmap depicting Pearson correlation coefficients between the expression levels of selected genes in advanced carotid lesions. Correlations with p < 0.01 are marked for statistical significance.

### Divergent Correlations of Kynurenine and NAD+ Pathway Genes with Macrophage Polarization

In the AE cohort, the shoulder and culprit lesion were dissected separately for transcriptomics and histology, potentially underestimating correlations. To address this, we used the MAASHPS biobank, where histology and RNA were derived from adjacent sections, allowing us to further investigate whether macrophage polarization (M1/M2) marker expression was associated with pathways involved in kynurenine formation and metabolism. The correlations between the expression of genes in the kynurenine pathway and markers of M1 and M2 macrophage polarization in the MAASHPS biobank are summarized in Table 2. In M2 marker macrophages, the expression of IDO1 (R = −0.49, p < 0.01), KYNU (R = −0.72, p < 0.001), and KMO (R = −0.60, p < 0.001) showed strong negative correlations, indicating lower expression levels in M2 macrophages. In contrast, HAAO expression was positively correlated (R = 0.48, p < 0.01). For the M1 marker positive macrophages, there were consistent, but no significant correlations in the opposite direction as for M2 marker positive macrophages. The association between higher *KMO* expression and M1 macrophages (R = 0.35, p < 0.05) was significant. Genes involved in de novo NAD+ synthesis, including *NADSYN1* and *NMNAT3*, showed a significant negative correlation with M2 macrophages (R = −0.49, p < 0.01 and R = 0.58, p < 0.001, respectively). Within the salvage pathway, *NAMPT* positively correlated with M1 markers and negatively with M2 markers.

**Table 2.** Correlations between differential expression of genes associated with NAD+ synthesis and macrophages positive for either M1 markers or M2 markers.

| Pathway |  | Gene | M1 macrophages |  | M2 macrophages |  |
| --- | --- | --- | --- | --- | --- | --- |
|  |  |  | R | p-value | R | p-value |
| Kynurenine |  | IDO1 | 0.29 | 0.11 | -0.49 | <0.01 |
|  |  | IDO2 | 0.05 | 0.81 | -0.26 | 0.16 |
|  |  | TDO2 | 0.17 | 0.37 | -0.47 | <0.01 |
|  |  | AFMID | 0.22 | 0.23 | -0.10 | 0.60 |
|  |  | KAT | 0.33 | 0.07 | 0.04 | 0.83 |
|  |  | KYNU | 0.34 | 0.06 | -0.72 | <0.001 |
|  |  | KMO | 0.35 | <0.05 | -0.60 | <0.001 |
|  |  | HAAO | -0.22 | 0.24 | 0.48 | <0.01 |
|  |  | ACMSD | -0.02 | 0.91 | 0.02 | 0.91 |
|  |  | QPRT | 0.07 | 0.71 | -0.08 | 0.65 |
|  | Preiss<br>Handler | NADSYN1 | 0.21 | 0.24 | -0.49 | <0.01 |
|  |  | NAPRT | 0.23 | 0.20 | -0.31 | 0.08 |
|  | Salvage | NMNAT1 | 0.28 | 0.13 | -0.23 | 0.21 |
|  |  | NMNAT2 | -0.19 | 0.30 | 0.02 | 0.93 |
|  |  | NMNAT3 | -0.12 | 0.51 | 0.58 | <0.001 |
|  |  | NAMPT | 0.39 | <0.05 | -0.53 | <0.01 |
|  |  | NRK1 | 0.03 | 0.87 | -0.02 | 0.91 |
|  |  | NRK2 | 0.10 | 0.59 | 0.08 | 0.68 |
|  | NAD+<br>consumption | CD38 | 0.15 | 0.43 | -0.42 | <0.05 |
|  |  | CD157 | 0.56 | <0.001 | -0.53 | <0.01 |

### In Vitro Validation Confirms Divergent Kynurenine and NAD+ Pathway Expression, Leading to Lower NAD+ in M1-Like Macrophages

We generated human monocyte-derived macrophages stimulated with either IFN-γ to resemble an M1-like phenotype or IL-4 for an M2-like phenotype. In line with our findings in the MAASHPS biobank, M2-like macrophages had significantly lower expression of the *IDO1*, *TDO2*, *CD38*, *CD157*, and *NAMPT* genes, but higher expression of *NMNAT3* compared to M1-like macrophages. We found no significant differences between both groups for *KYNU*, *HAAO*, and *NADSYN1*. Contradictory to the MAASHPS biobank, we found a significant decreased expression of *QPRT* in M1-like macrophages but higher expression of *KMO* in M2-like macrophages.

Furthermore, at the metabolite level, tryptophan levels were significantly lower in M1-like macrophages compared control (*p* < 0.05) to M2-like macrophages (*p* < 0.05). We also observed significantly lower levels of NAD+ in M1-like macrophages compared to the control group (*p* = 0.001) (Figure 4E). NADH showed a similar trend to NAD+ with significantly lower levels in the M1-like macrophages compared to NADH+ (*p* < 0.001), but we found no difference between the NADH/NAD+-ratio.

**Figure 4.**
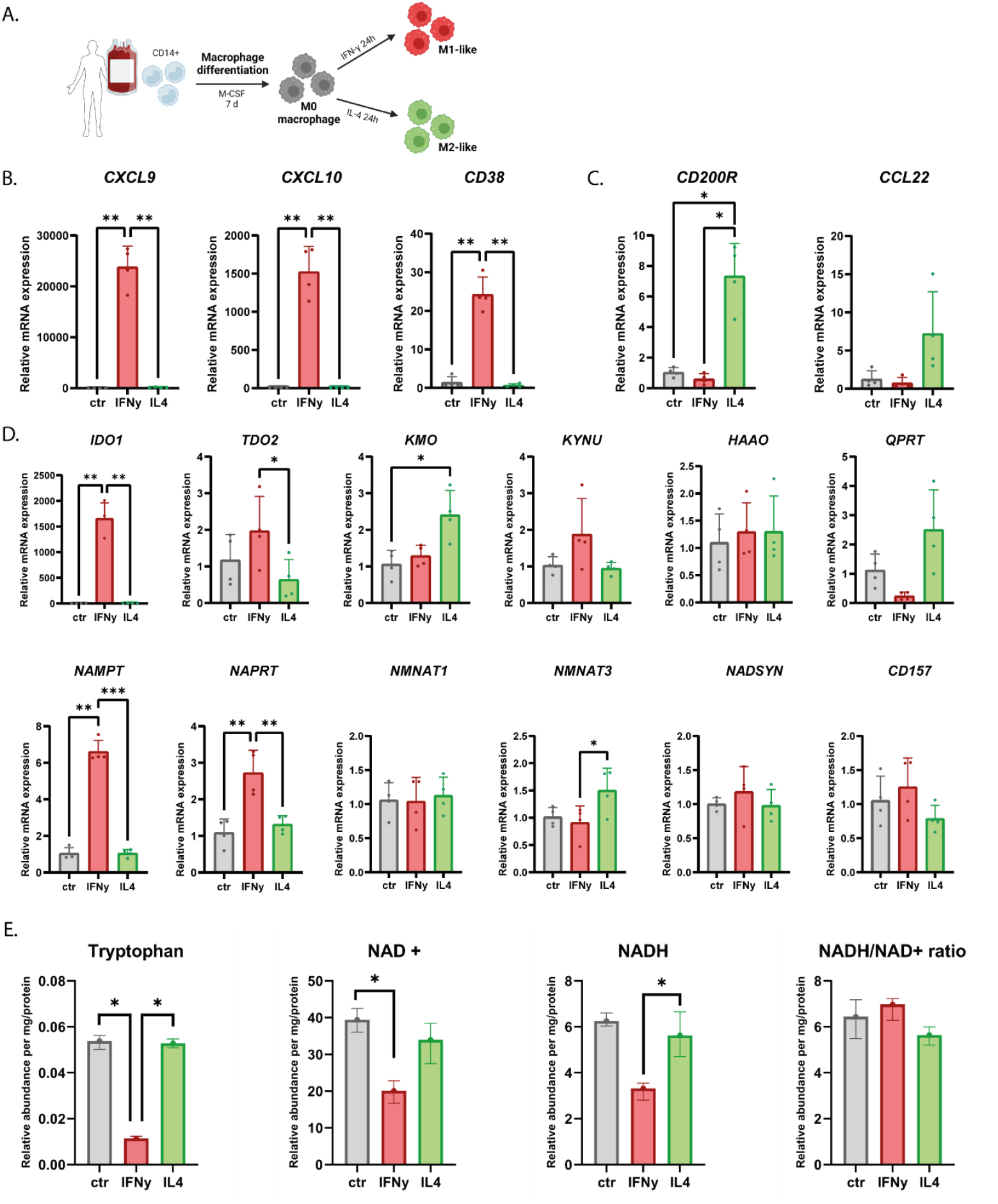
Differences in expression of kynurenine pathway enzymes observed in vivo are also replicated in in vitro M1- and M2-like macrophages. A. Human monocytes were differentiated according to the protocol and stimulated with IFNγ or IL4 for 24 hours. Relative normalized mRNA expression of M1-like (B) and M2-like (C) genes. D. Relative normalized mRNA expression of genes in the kynurenine pathway. E. Relative abundances of Trp, kynurenine, NAD+ and the NAD+/NADH-ratio P<0,05 = *, P<0.01 = **, P<0.001=***.

## Discussion

Our study identified two key findings that shed light on the importance of the kynurenine pathway in determining macrophage phenotype and atherosclerosis progression. First, transcriptomic data from the MAASHPS biobank revealed that the expression of key genes involved in the kynurenine pathway, such as *IDO1*, *KYNU*, *KMO*, and *HAAO*, were negatively correlated with M2-like macrophages, indicating reduced kynurenine pathway activity. In contrast, M1 macrophages demonstrated positive associations of these enzymes, highlighting their specific role in kynurenine production, as attested by our in-vitro analysis.

Second, we found that M1-like macrophages in vitro demonstrated increased expression of *IDO1* with a concomitant reduced expression of *QPRT*; the enzyme responsible for converting quinolinic acid into NAD+. Concentrations of NAD+ were also lower in M1-like macrophages. This dual effect suggests that while IDO1 drives upstream kynurenine production, QPRT downregulation impairs NAD+ synthesis, potentially contributing to the inflammatory phenotype in macrophages. Together, these results highlight the distinct regulation of the kynurenine pathway in M1-like macrophage subtypes and suggest that increased concentrations of kynurenine are not merely a reflection of low grade inflammation, but a specific reflection of inflammatory M1-like macrophage activity.

Increased concentrations of kynurenine have been associated with a greater risk of cardiovascular disease (CVD) and mortality (7). Previous studies have predominantly focused on kynurenine as a ‘passive’ biomarker. Increased concentrations of kynurenine were merely seen as a reflection upregulated IDO1 activity. Increased IDO1 activity increased kynurenine plasma concentrations, primarily due to cytokines such as IFN-γ and IL-6 in the context of low-grade inflammation (31). Our findings challenge this view. In the AE biobank, we found no increase in the kynurenine-to-tryptophan ratio, indicating no increase in the conversation rate of tryptophan into kynurenine via IDO. The lack of increased *IDO1* expression despite elevated macrophage numbers in plaques in this cohort also suggest that CD68+ macrophages are not automatically associated with increased levels IDO1 activity. However, the transcriptomic data in the MAASHPS biobank support the concept that kynurenine accumulation must reflect specific M1-like macrophage activity, since we found a significant negative correlation between M2-like phenotype macrophages and *IDO1* expression in this cohort.

Our in vitro experiments demonstrate that this accumulation might be mediated by a distinct *QPRT* downregulation in M1-like macrophages. This impaired downstream conversion towards NAD+, coupled with increased upstream kynurenine production, provides a plausible explanation for the trend in elevated kynurenine levels observed in plaques with higher macrophage content. However, one of the main limitations is the lack of macrophage subtype-specific data in the AE biobank, likely explaining the absence of significant outcomes across macrophage groups in this cohort.

We thus demonstrate that elevated kynurenine concentrations could specifically reflect M1-like macrophage activity due to specific metabolic changes within these subtypes of macrophages, and the total macrophage content of atherosclerotic plaques is insufficient to capture nuanced alterations in the kynurenine pathway. This underscores the need for macrophage subtype-specific analyses to better understand the role of macrophage kynurenine metabolism in atherosclerosis.

Kynurenine metabolism could potentially be used as a signaling molecule by M1 like macrophages. Multiple metabolites downstream of IDO1 and upstream of QPRT are a ligand for the aryl hydrocarbon receptor (AhR), a pathway implicated in diverse immunological processes in atherosclerosis (32, 33). Kynurenine’s ability to act as a signaling molecule may influence intercellular communication, particularly given its capacity to cross the cell membrane (32). AhR expression in macrophages and T-cells (34) further underscores the kynurenine pathway’s dual role as both a signaling and metabolic axis in atherosclerosis. On the other hand, *QPRT* downregulation also results lower NAD+ levels, an essential cofactor for energy metabolism. Both these phenomena, kynurenine-AhR signaling as well as NAD+ depletion may contribute to an inflammatory state in influence and macrophage metabolism under inflammatory conditions (35). The reduction in NAD+ likely drives macrophages toward an inflammatory, glycolytic phenotype, consistent with prior findings (21). These metabolic changes promote plaque instability and may exacerbate cardiovascular risk (36). Notably, we observed no significant differences in the NADH/NAD+ ratio.

Our findings also align with recent work in cancer biology, where QPRT inhibition in M2-like tumor-associated macrophages (TAMs), reduces immune evasion and promotes inflammatory responses (37). In cancer, M2-like macrophages promote tumor growth as they orchestrate immune evasion and suppression of inflammation (38, 39). Inhibiting QPRT disrupts this process and restores inflammation, which is crucial to halt tumor growth in the context of cancer therapy (40, 41). By contrast, in atherosclerosis, inflammatory M1 macrophages drive pathology, and their downregulation of QPRT sustains inflammation. This context-specific role highlights the importance of understanding the function of QPRT across macrophage subtypes and different disease settings.

Although QPRT modulation offers potential therapeutic benefits, its wide expression in multiple human tissues, including the brain, liver, intestine, and kidneys, poses feasibility challenges. QPRT knockout models in mice have demonstrated significant physiological consequences, such as decreased neuromuscular function and increased frailty, when QPRT is deleted and mice were no supplement with a different precursor of NAD + other than TRP (42).

Instead, rather than targeting QPRT directly, bypassing de novo NAD+ synthesis altogether through NAD+ precursor supplementation (e.g., nicotinamide riboside or nicotinamide) emerges as a more viable strategy. Supplementation could restore NAD+ levels, mitigate macrophage-driven inflammation, and promote plaque stability. However, the results of supplementing NR in a different context than atherosclerosis so far has led to mixed results in humans. Despite the fact that it has been shown that NR intake does increase serum NAD+ (22). Furthermore, it is import to consider that this approach would not directly address kynurenine accumulation and therefore also would address the intracellular communication via the AhR between macrophages and T-cells.

## Conclusion and Future Directions

Our study challenges the traditional view that IDO1 alone drives kynurenine accumulation in atherosclerosis. Instead, we identify a potential role for QPRT downregulation as a metabolic bottleneck linking disrupted NAD+ synthesis to M1-like macrophage polarization. While our findings demonstrate an association between elevated kynurenine levels and M1-like macrophage activity, they do not establish a causal relationship. Rather, these results suggest that kynurenine accumulation may serve as a marker of macrophage-driven metabolic dysregulation rather than a direct driver of inflammation.

This dual disruption, upstream kynurenine accumulation and downstream NAD+ depletion, may contribute to a pro-inflammatory, metabolically dysregulated environment in atherosclerotic plaques. Given this metabolic interplay, targeting NAD+ precursor supplementation, rather than QPRT itself, could represent a potential strategy to restore NAD+ levels and mitigate M1-driven inflammation.

Future studies should explore whether therapeutic modulation of the kynurenine pathway or NAD+ supplementation could influence macrophage polarization and, ultimately, cardiovascular risk in patients with atherosclerosis.

## Supplements

**Suplementary table 1.**
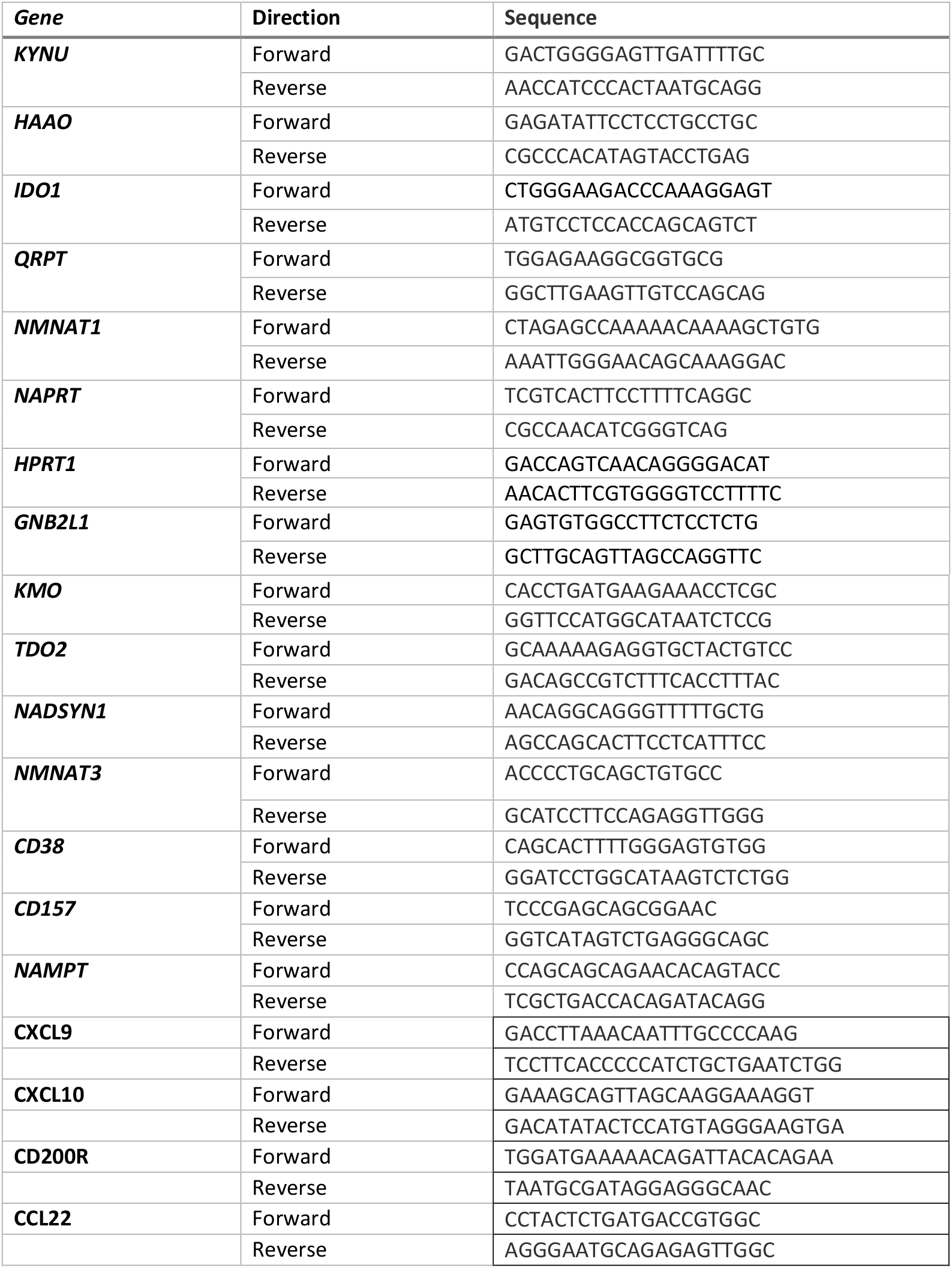
Primer sequences.

